# PyZebraScope: an open-source platform for brain-wide neural activity imaging in zebrafish

**DOI:** 10.1101/2022.02.13.480249

**Authors:** Rani Barbara, Madhu Nagathihalli Kantharaju, Ravid Haruvi, Kyle Harrington, Takashi Kawashima

## Abstract

Understanding how neurons interact across the brain to control animal behaviors is one of the central goals in neuroscience. Recent developments in fluorescent microscopy and genetically-encoded calcium indicators led to the establishment of whole-brain imaging methods in zebrafish, which records neural activity across a brain-wide volume with single-cell resolution. Pioneering studies of whole-brain imaging used custom light-sheet microscopes, and their operation relied on commercially developed and maintained software that is not available globally. Hence it has been challenging to disseminate and develop the technology in the research community. Here, we present PyZebrascope, an open-source Python platform designed for neural activity imaging in zebrafish using light-sheet microscopy. PyZebrascope has intuitive user interfaces and implements essential features for whole-brain imaging, such as two orthogonal excitation beams and eye damage prevention. Its modular architecture allows the inclusion of advanced algorithms for microscope control and image processing. As a proof of concept, we implemented an automatic algorithm for maximizing the image resolution in the brain by precisely aligning the excitation beams to the image focal plane. PyZebrascope enables whole-brain neural activity imaging in fish behaving in a virtual reality environment with a stable high data throughput and low CPU and memory consumption. Thus, PyZebrascope will help disseminate and develop light-sheet microscopy techniques in the neuroscience community and advance our understanding of whole-brain neural dynamics during animal behaviors.

## Introduction

Animal behaviors occur through the collective actions of neurons distributed across the brain. Understanding such distributed neural dynamics in their entirety has been one of the central goals in neuroscience^1^. Toward this goal, optical recording of neural activity at a brain-wide scale has become possible based on recent developments in genetically-encoded calcium indicators^2,3^ and volumetric fluorescence microscopy^4–7^. Whole-brain neural activity imaging at cellular resolution was first demonstrated in larval zebrafish^8–10^ among other vertebrate model organisms using digital scanned laser light-sheet microscopy (DSLM)^4^. DSLM excites sample fluorescence in multiple voxels along the light cone of the excitation beam, and rapid scanning of the excitation beam enables fast volumetric scans with high spatial resolution and low light toxicity. These advantages of DSLM are best exploited in the optically transparent brain of larval zebrafish. DSLM enabled studies of whole-brain neural dynamics during visually-evoked swimmming^11,12^, motor learning^13,14^, learned helplessness^15^, threat escape^16^, and body posture change^17^. Usage of the DSLM also revealed spontaneous noise dynamics across the brain^18^ and how those dynamics change during neural perturbations^19,20^ or administration of psychoactive reagents^21^. DSLM also enables high-speed voltage imaging on a single axial plane for recording membrane potential and spiking activity from a neural population during swimming in the midbrain^22^ and the spinal cord^23^. These pioneering studies in zebrafish using light-sheet microscopy expanded our understanding of diverse functionalities of the vertebrate brain.

Despite its advantages, it is still challenging to build a light-sheet microscope customized for zebrafish imaging and operate it for multiple experiments in a day. These challenges come from the complexity of the microscope itself and its built-in parameters that the experimenter needs to manipulate. For example, light-sheet microscopes for whole-brain imaging in zebrafish (**Fig. 1a**) typically consist of two excitation optical paths from the lateral and front sides of the fish and one optical detection path above the fish ^11,14,16,17^ (**Fig. 1b**). The fish is fixed in a water chamber and needs to be precisely maneuvered into focal points of the excitation and detection objectives. Moreover, it is necessary to prevent laser illumination into the eyes to secure fish’s vision for behavioral tasks and prevent eye damage (**Fig. 1b**). This configuration requires the experimenters to set at least ∼20 parameters (camera exposure time per plane, number of planes per volume, the start and end positions for 2d motion for each excitation beam, light intensity and on/off timing for lasers, the start and end positions for detection objective motions, 3-dimensional positions for the fish chamber, and parameters for eye exclusion). The first studies^8,11^ on whole-brain imaging in zebrafish were made possible by using software custom-developed and maintained by a commercial entity, which charges high-priced service costs and does not provide service globally. This situation prevented the dissemination of the technology in a flexible and customizable manner. Past progress of optical microscopy in neuroscience has been driven by open-source software^24,25^ for microscope control written in a programming language widely used in academics, such as ScanImage^24^ for multiphoton microscopy. Hence, developing an open-source platform for light-sheet microscopy dedicated to neural activity imaging in zebrafish is necessary. Such software will reduce the laboriousness of imaging experiments by providing control of numerous device parameters on a simple and intuitive interface. Its implementation in a standard programming language for researchers will facilitate the addition of innovative functionalities toward their research goals by importing modules from a broad developer ecosystem for hardware control, signal processing, computer vision, and machine learning.

**Figure 1:**
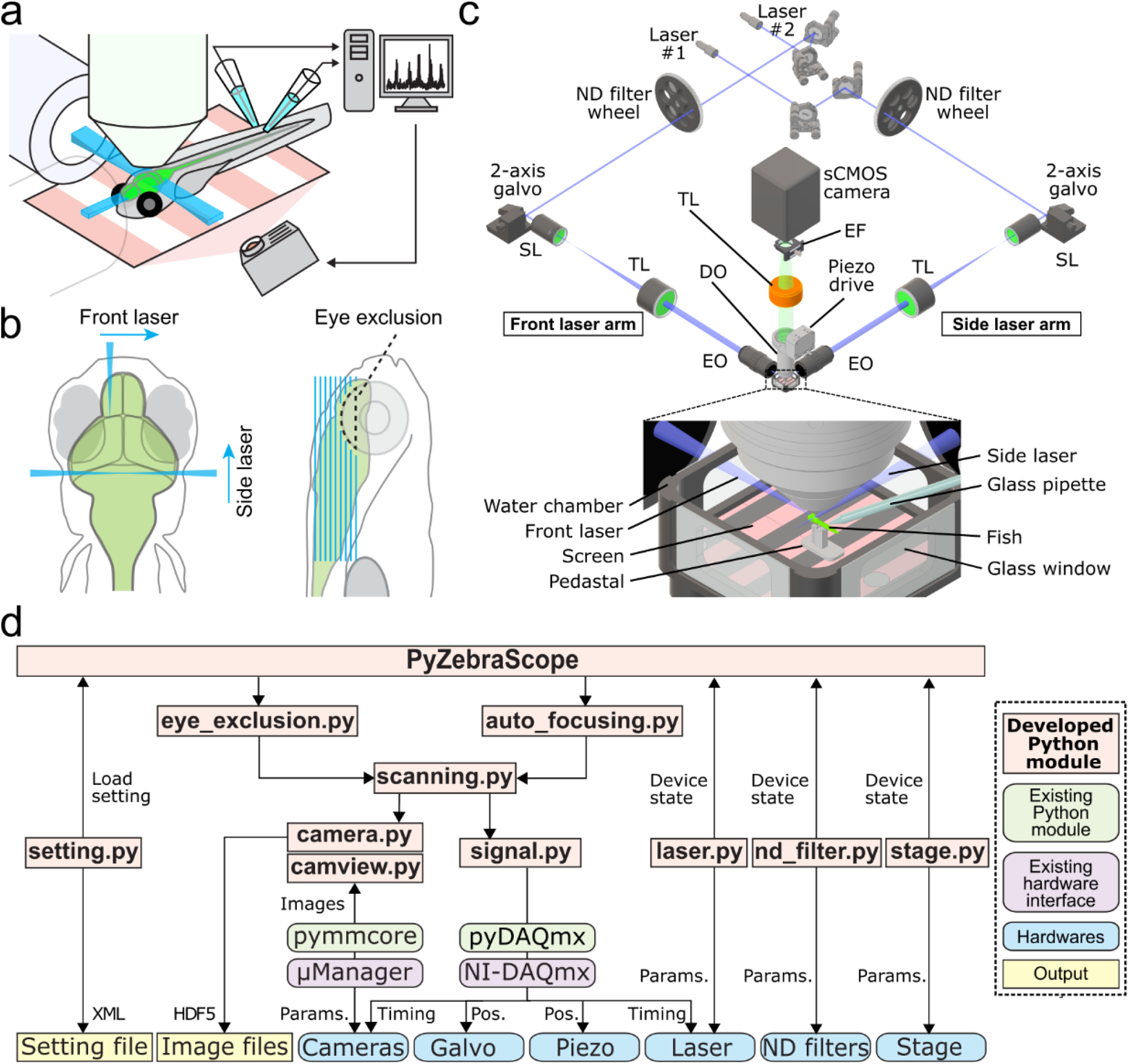
Our microscope setup and modular architecture of PyZebrascope for whole-brain neural activity imaging in zebrafish. **(a)** Schematic drawing of setups for whole-brain neural activity imaging in behaving zebrafish. Excitation objectives on the lateral and front sides of the fish illuminate excitation beams into the brain. The detection objective above the fish moves in the axial direction in sync with the motion of the excitation beams during the volumetric scan. We paralyze the muscle of the fish and record swim-related electrical signals from axonal bundles of spinal motoneurons by using two pipettes attached to the tail. The recorded signals are analyzed online and reflected in the motion of visual scenes projected below the fish. **(b)** Two-beam scanning during whole-brain imaging. The front laser beam scans areas between the eyes, and the side laser beam scans areas behind and above the eye. The side laser beam avoids eye damage by turning off when it overlaps with the eye. **(c)** Schematic of our light-sheet microscope and water chamber for imaging zebrafish. Detailed descriptions are in the main text. TL, tube lens; SL, scan lens; EO, excitation objective; DO, detection objective; EF, emission filter. **(d)** The architecture of PyZebrascope software. Our developed modules (beige), existing Python libraries (green), hardware drivers (purple), hardware (blue) and output files (yellow) are shown. Arrows show information flow between modules with labels such as “Pos.” (linear position information for devices) and “Params.” (device parameters).

Toward this goal, we developed PyZebrascope, an open-source Python platform designed for whole-brain imaging in zebrafish using light-sheet microscopy. PyZebrascope implements essential features for whole-brain imaging, such as two orthogonal excitation beams and eye damage prevention. Its user interfaces allow the users to adjust parameters for lasers, filters, beam scanning, camera, sample positions, and eye damage prevention intuitively. It is also possible to add advanced algorithms for microscope control and image processing developed in the Python community^26,27^. As a proof of concept, we implemented a GPU-based automatic algorithm for maximizing the image resolution in the brain by precisely aligning the excitation beams to the image focal plane, which is usually a time-consuming and indecisive process for the experimenter. Lastly, we demonstrated that PyZebrascope enables whole-brain imaging in zebrafish behaving in a virtual reality environment with stable high data throughput and low CPU and memory consumption. Thus, PyZebrascope is a versatile platform for disseminating and advancing technology for large-scale neural activity imaging in zebrafish and will accelerate our understanding of whole-brain neural dynamics during animal behaviors.

## Results

### Microscope design and the architecture of PyZebrascope

We developed PyZebrascope for our custom-built light-sheet microscope (**Fig. 1c and S1, Supplementary Table 1** for parts lists), for which many components are common to those in the previous studies^9,11,14,16^. Our system is equipped with two lasers, providing light sources for two excitation arms that are required to scan fish brains from the lateral and front sides **(Fig. 1b**). We control the lasers’ on/off by digital input into the lasers. The brightnesses of laser outputs are controlled by (i) setting the power level of the laser using serial communication and (ii) diminishing the laser output with different levels of neutral density (ND) filters. These two levels of brightness adjustment allow us to maintain laser output levels in a range that has less power fluctuation. Typically, the excitation beam for the front scanning (**Fig. 1b**) requires less output power due to its narrow scanning range. Each brightness-adjusted beam is then scanned by sets of 2-axis galvanometers for volumetric scanning. The beams expand through a pair of a telecentric scan lens (SL) and a tube lens (TL) and then focus onto the sample through an excitation objective (EO). Fluorescence from the fish brain is collected by a detection objective (DO) placed above the brain, and this detection objective moves along the axial direction in sync with the excitation beam by a piezoelectric drive during volumetric scanning. The fluorescent image is focused onto scientific CMOS (sCMOS) camera through a tube lens (TL) and an emission filter (EF, **Fig. 1c**). We can also add another sCMOS camera in the detection path for multicolor imaging, as shown in our CAD model (**Fig. S1**, CAD model files available on request).

PyZebrascope controls the above-mentioned multiple devices through its modular architecture and organized user interface (**Fig. 1d, 2 and S2**). It does not require compiling and is launchable from any Python development environment that allows PyQT applications. It controls devices through serial communications (laser setting, filter wheels, sample stage), analog output (galvanometer, camera timing), digital output (laser on/off), and camera interface cables (**Fig. S2**). The camera unit (camera.py) controls camera settings through a Python binding (pymmcore, https://github.com/micro-manager/pymmcore) for low-level device interface of μManager^28^. It also receives acquired images from the camera and saves them in HDF files. The analog/digital output unit (signal.py) generates waveforms and outputs them through a Python binding (ni-daqmx, https://nidaqmx-python.readthedocs.io/) for data acquisition interface devices. Its outputs can be modified to drive alternative components of light-sheet microscopes that operate according to voltage inputs, such as electrically tunable lens^29^ and scanners based on microelectromechanical systems (MEMS)^17^. Other modules for controlling devices via serial communication (laser.py, nd_filter.py, piezo.py) are written in a generalizable manner so that external users can adapt them to their preferred devices.

The configuration unit (setting.py) allows the users to save all the microscope configuration parameters in an XML file. Savable parameters include the state of all connected devices, the position of the 3-axis stage, the choice of lasers and their output intensity, type of neutral density filters, the region of interest on the camera’s field of view, and parameters for volumetric scanning. This file is also automatically saved for every experiment with time logs. The experimenter can load this setting file later to connect necessary devices and set all the device parameters automatically. Although the user still needs to fine-tune sample positions and microscope scanning parameters for each experiment, this automatic loading minimizes the users’ efforts in conducting experiments consistently across sessions.

### Sample preparation and stage control

The fish for imaging resides in a water chamber with glass windows (**Fig. 1c**). Within the water chamber, the fish is embedded in low-melting agarose on a small pedestal, and this pedestal allows us to expose the fish head after removing surrounding agarose and at the same time record electrical signals from the tail for fictive swim recording^11,13,30^. The user mounts the fish into the water chamber outside the microscope at the beginning of each experiment. Then the chamber is placed on a motorized 3-axis stage, which brings the fish to the focus of the detection and excitation objectives (**Fig. S1**).

The PyZebrascope sample stage unit (stage.py) moves the motorized stage holding the water chamber according to the user’s commands from the software interface. It also allows users to automatically move the stage between stereotypical positions, such as a home position for replacing water chambers and an imaging position for performing experiments. During imaging experiments, the water chamber, the detection objective, and the excitation objectives need to be placed in close physical proximity. Therefore, the stage unit moves the stage in a constrained manner to avoid the risk of accidental collisions between these components.

### User interface and waveform outputs of Pyzebrascope

We designed graphical user interfaces based on QT Designer (**Fig. 2**, https://doc.qt.io/qt-5/qtdesigner-manual.html). All of the above device parameters are organized into two tabs in the main window (**Fig. 2a**). Waveforms of analog and digital outputs (**Fig. 2b**) can be viewed in the user interface window. We implemented a waveform-generating algorithm that synchronizes the beam scanning to the full opening of the rolling shutter in the sCMOS camera and the movement delay of the piezoelectric drive of the detection objective. It moves the piezoelectric drives and the beam scanning in the axial direction in a continuous manner (**Fig. 2b**) rather than in a step-wise manner because the piezoelectric drive cannot move the weight of the detection objective with sub-millisecond accuracy. Its continuous motion without frequent accelerating/decelerating events provides a better match of axial positions between the detection objective and beam scanning. This algorithm uses Numpy^26^, a common library for array programming for researchers, and can be flexibly edited in the code (signal.py) to enable different types of scanning patterns such as bidirectional scanning for faster volumetric imaging^31^.

**Figure 2:**
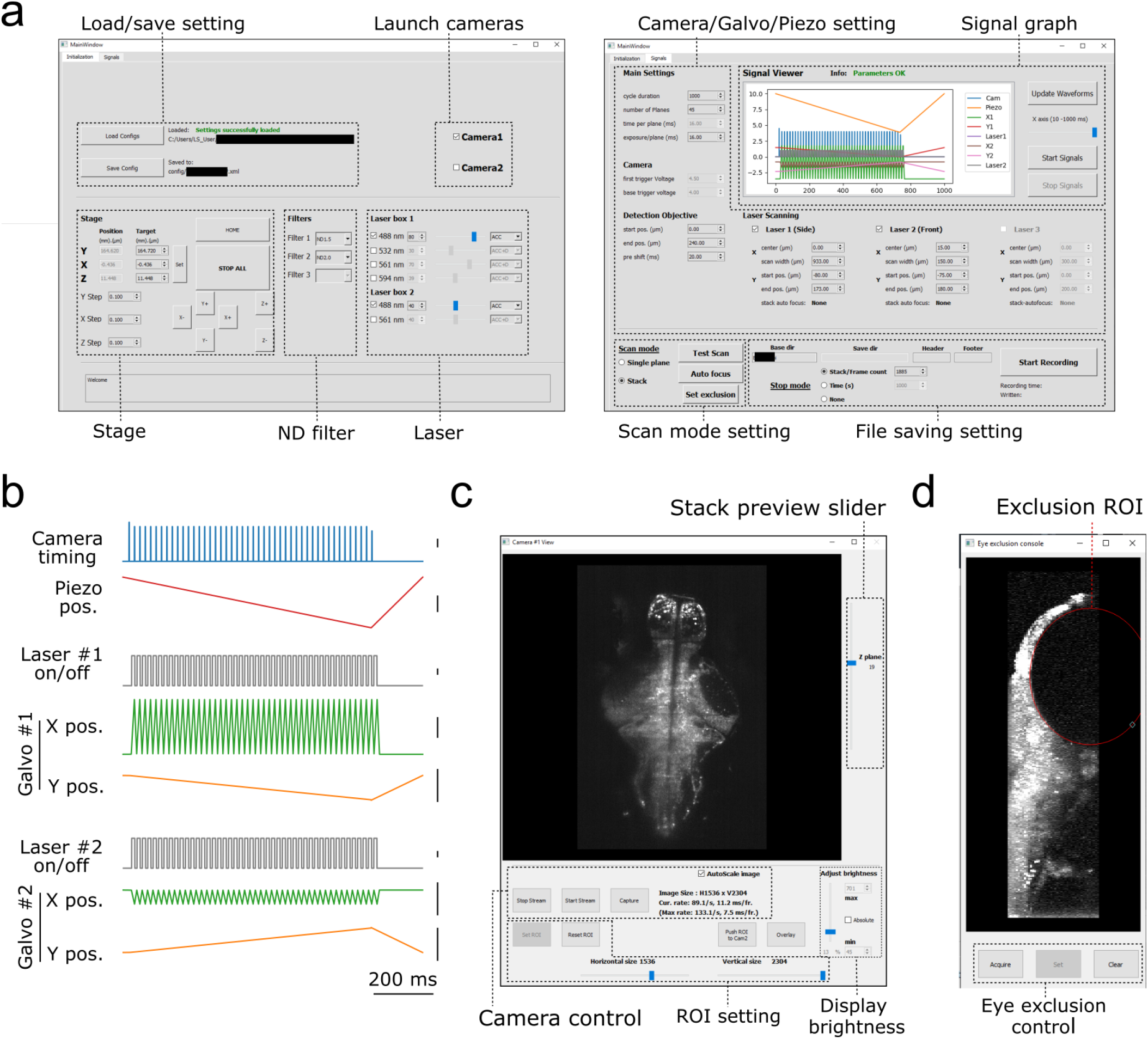
Graphical user interface of PyZebrascope. **(a)** Two main tabbed windows of the user interface of PyZebrascope. The first window provides saving/loading microscope configurations, sample stage control, selection of neutral density filter, and laser control. The second window provides camera exposure control, motion control for the detection objective, scanning control for excitation beams, switch between volumetric and single-plane scans, and file logging control. **(b)** Examples of analog and digital outputs for microscope components during a volumetric scan. Black scale bars next to waveforms represent 1 V. Camera trigger signals have different amplitudes at the start (4.5 V) and the end of a volumetric scan (3.5 V) compared to those in the middle (4.0 V). These varying amplitudes of trigger signals above the camera’s threshold would allow the behavior control software to receive camera trigger signals in parallel and detect the onset of each volumetric scan for synchronization. **(c)** Camera view window. It allows users to view ambient and fluorescent images, scroll through different Z-planes of a volumetric stack, zoom in/out, adjust the image display brightness, set region of interest (ROI) to the camera, and overlay two images in different color channels from two cameras during multicolor imaging. This window can also be stretched to any preferred size. **(d)** Eye exclusion window. The lateral view of a volumetric stack allows the user to set an elliptic exclusion area (red) which will turn off the side laser when it scans over the eye.

The camera view window (**Fig. 2c**, camview.py) will pop up separately when the camera is turned on. It allows the users to view the ambient and fluorescent images of the sample and adjust the display brightness and zoom. It also allows the inspection of the quality of the volumetric stack by using a slider for changing the displayed Z planes. PyZebrascope supports two cameras in the detection path for multicolor imaging, and this camera view window can display images from two cameras separately or overlay them in RGB color channels.

PyZebrascope also has a dedicated module and interface for eye damage prevention (eye_exclusion.py) (**Fig. 2d**), where the experimenter looks at the side projection of a volumetric stack and draws an elliptic region of interest (ROI) to set where the lateral laser should be turned off (**Fig. 1b**). This module automatically calculates the timing for turning off the laser across multiple Z-planes based on the set ROI. Thus, this interface prevents laser illumination into the eye while performing experiments, enabling testing of behavioral tasks that depends on visual features presented to the fish.

### Automatic alignment of excitation beams to image focal planes

The Python software ecosystem offers a variety of highly optimized libraries for image processing and machine learning. Such distinct advantages for using Python, as well as the modular architecture of PyZebrascope, enable the implementation of advanced algorithms for microscope control and online image analyses. As a proof of concept to demonstrate this advantage, we developed an autofocusing module (auto_focusing.py) that adjusts the axial position of the excitation beam in the sample to the best focus of the detection objective (**Fig. 3a**). This procedure is usually a time-consuming manual step during the preparation of whole-brain volumetric imaging for each fish. The experimenter needs to align the position of the excitation beam from the superficial part to the deep part of the brain. This alignment is necessary for both the front and side excitation beams. In addition, images at the bottom part of the brain are typically blurry due to the diffraction of fluorescence through the sample, which makes the alignment indecisive for the user.

**Figure 3:**
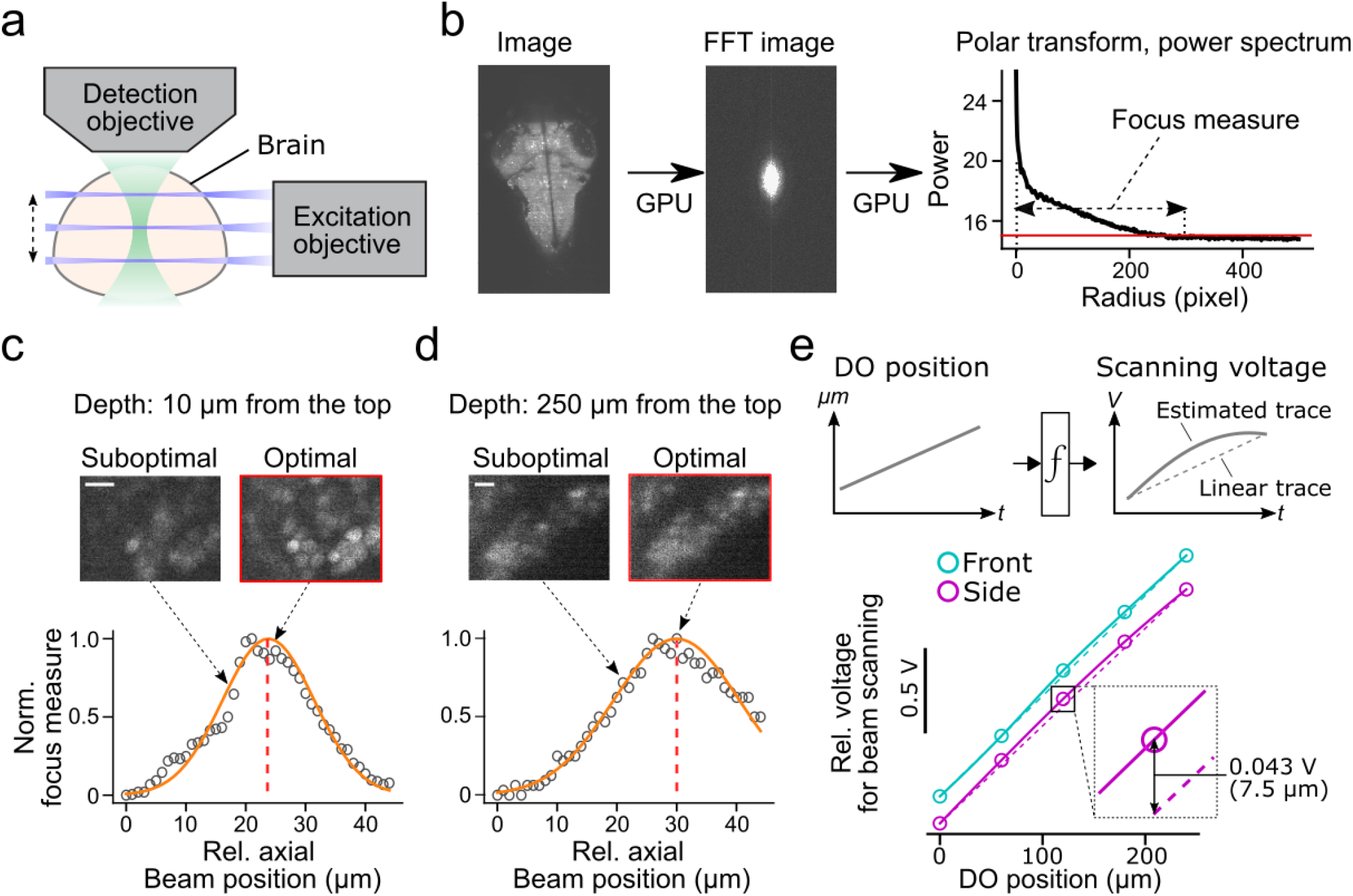
Automatic alignment of excitation beams to the image focal plane. **(a)** Schematic of the alignment between the excitation beams and the detection objective based on acquired fluorescent images in the brain. **(b)** Calculation of focus measures. We apply a Fourier transform to the image, followed by a polar transform to the resulting power spectrum and a 1D projection along the angular axis. Then we calculate the width of the power spectrum above the threshold as a focus measure. Images with the finest resolution yield a higher focus measure. **(c)** Detection of the optimal beam position for the dorsal part of the brain. Normalized focus measures at different beam positions (circles), Gaussian fit (orange line), and the best focus (red dashed line) are plotted. Sample images from different beam positions are shown on the top. Scale bar, 10 μm. **(d)** Same plots as (c) for the ventral part of the brain. Scale bar, 10 μm. **(e)** Top, estimation of the nonlinear function between the position of the detection objective (DO) and analog output voltage for the axial scanning of the excitation beams during a volumetric scan. The estimated function (solid lines) may show nonlinearity compared to linear estimation based on the start and end position of the detection objective (dotted lines). *Bottom*, we estimated nonlinear functions between the positions of the detection objective (DO) and analog output voltage for axial beam scanning in the brain of a zebrafish. We determined optimal voltage output at five positions of the detection objective during a volumetric scan of the front beam (cyan) and side beam (magenta) independently and estimated transformation functions by cubic interpolation. The resulting functions (solid lines) showed nonlinearity compared to linear estimation (dashed lines). *Bottom inset*, the difference between the optimal analog output and the linear estimation was as large as 0.043V, which amounts to 7.5 μm of the axial position of the side beam, at the middle plane of a volumetric scan.

We implemented an automatic algorithm to assist such a time-consuming alignment process by using an image resolution measure based on Fourier transformation^32^ (**Fig. 3b**). Optimal beam alignment results in higher resolution of fluorescent images, which corresponds to higher powers in the high-frequency domain in Fourier-transformed images. This algorithm first Fourier-transforms the fluorescent image to obtain its 2D power spectrum. Polar transform is applied to the 2D power spectrum and is projected onto 1D by averaging along the angular dimension. We then use the logarithm of the obtained 1D projected array to quantify image resolution. The threshold for quantifying image resolution is defined as the sum of mean and three times the standard deviation of the baseline part of the power spectrum. We then count the number of points above the set threshold and use this number as an image resolution measure (**Fig. 3b**). We implemented this algorithm using a library for GPU computing (CuPy, https://cupy.dev/) to minimize computation time for Fourier transform and polar transform.

Our auto-focusing module searches for the best focus by acquiring images at different axial beam positions (41 planes at an interval of 1 μm) for a given position of the detection objective (DO). It calculates the above resolution measure for each acquired plane and normalizes resolution measures to between 0 (poor resolution) and 1 (best resolution) across different planes. The peak position was detected by fitting a Gaussian distribution function to the normalized resolution measures, and the center of the estimated distribution was designated as the best focal plane. These sampling and computing processes take less than 2 seconds in total and accurately detect the best focus for the dorsal (**Fig. 3c**) and ventral (**Fig. 3d**) part of the fish brain. We further applied this technique to estimate the nonlinear function between the position of the detection objective (DO) and voltage commands for axial positioning of the excitation beams during a volumetric scan (**Fig. 3e**). Such nonlinearity occurs because the angle of the scanning galvanometer is not necessarily linear to the axial position of the excitation beam. Our auto-focusing module automatically finds the best axial positions of the excitation beams, independently for the side and front beams, for five different DO positions along the Z-axis of a volumetric scan. It then estimates the optimal transformation function between the DO movements and scanning voltage for the excitation beams. Real transformation functions obtained in a zebrafish brain showed nonlinearity (**Fig. 3e**), and the optimal beam position at the middle of a volumetric scan differed from the linear estimation by 7.5 μm, a large enough distance to affect image resolution significantly (**Fig. 3c and d**). Thus, this algorithm allows us to obtain accurate alignment between the objective and the excitation beam beyond manual adjustments based on the start and end position of the volumetric acquisition.

### Whole-brain imaging in behaving zebrafish

We tested whether PyZebrascope can perform whole-brain imaging in zebrafish while recording its behavior during a behavioral task (**Fig. 4a**). We used a 6-day old transgenic zebrafish that pan-neuronally express nuclear-localized, genetically-encoded calcium indicators^33^ (HuC:H2B-GCaMP7f). The tested fish was able to react to the visual stimuli presented from below (**Fig. 4b**) and performed the motor learning task described in our previous work^13^. We were able to acquire continuous volumetric scans (45 planes at 1Hz) across the entire brain (**Fig. 4a**), from the extremities of the dorsal to the ventral regions (cerebellum and hypothalamus, respectively) and from extremities of the rostral to the caudal regions (forebrain and area postrema, respectively). The data rate was 300 MB per second, and the software was stable for the entire experiment duration (30 minutes). The CPU and memory usage stayed below 10% and 3 GB, respectively, and we did not experience any writing delays (see Methods for microscope control computer specifications).

**Figure 4:**
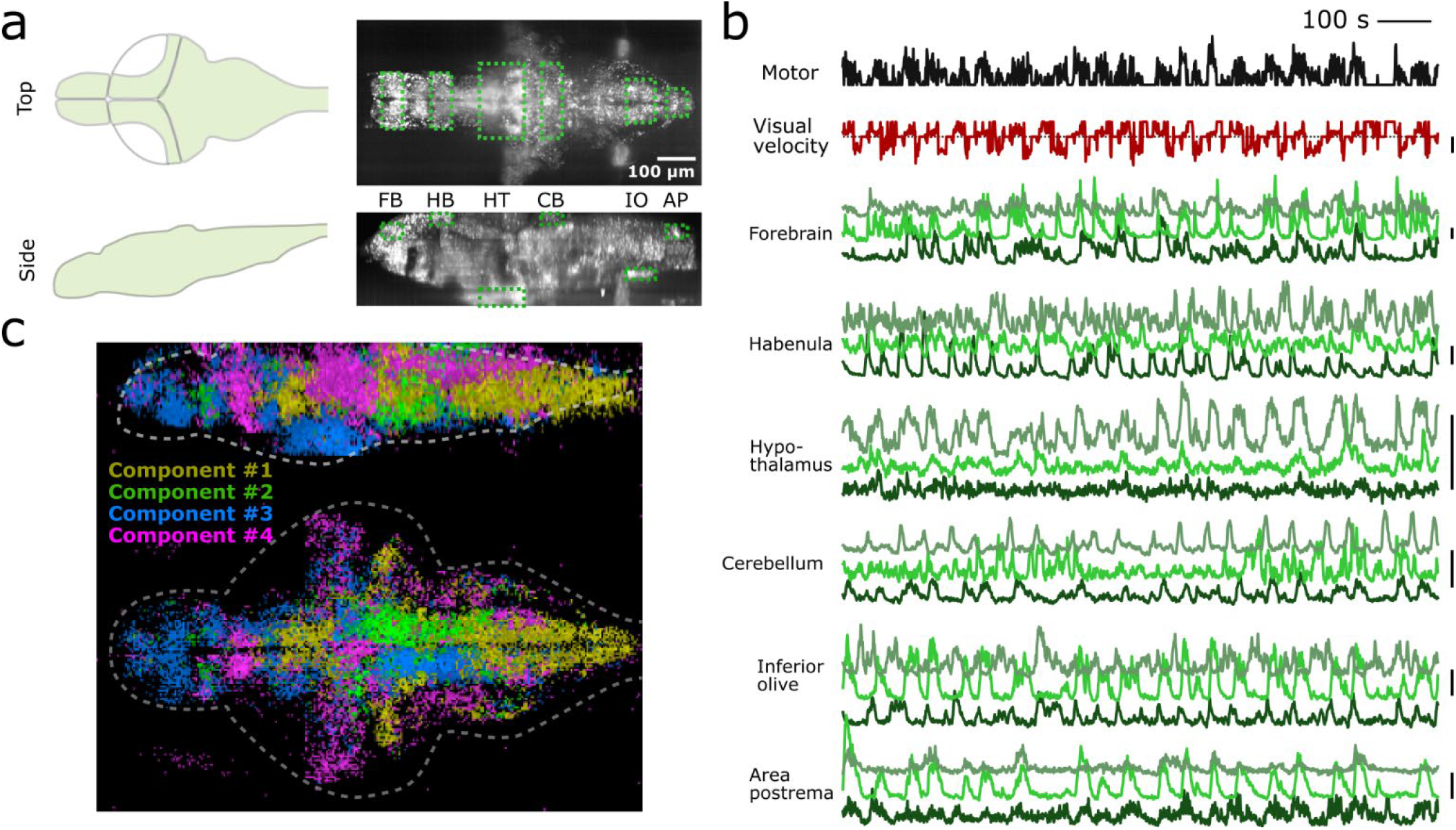
PyZebrascope enables stable recording of neural activity at a brain-wide scale in behaving zebrafish. **(a)** Scanning area of whole-brain imaging. The scan area covers the most dorsal part of the brain (cerebellum = CB, habenula = HB), the most ventral part of the brain (hypothalamus = HT), the most rostral part of the brain (forebrain = FB), and the most caudal part of the brain (Area postrema = AP, inferior olive = IO). **(b)** Simultaneous recording of swimming events (black), the visual velocity of the environment (red), and neural activity traces (green) of representative neurons from brain areas designated in (a) during a motor learning task. Three neurons are selected from three groups that show distinct activity patterns from each area based on k-means clustering methods. Scale bars on the right represent 100 % change in ΔF/F. Scale bar for visual velocity represents 2 mm/s, and traces below dotted lines represent backward motions of the environment triggered by swimming. **(c)** Whole-brain spatial map of neural activity clusters classified based on non-negative matrix factorization (n=4) of 36,168 neurons. Detailed descriptions are in the main text and method section.

We analyzed the acquired data by using image registration and segmentation algorithms developed in our previous work^13^. We identified ∼80,000 neurons across the brain, and their neural activity patterns remained stable for the entire duration of the experiment from the above areas of the brain (**Fig. 4b**). These results demonstrate PyZebrascope’s robustness for continuously acquiring large volumetric data over time. We tested the reliability of the data acquired by PyZebrascope by examining whether it is possible to identify behavior-related neural ensembles identified in other studies. We applied non-negative matrix factorization (4 set components) to the neural activity of all neurons across the entire brain and mapped the spatial locations of neurons that have significant weights for the identified components (**Fig. 4c**). We were able to identify a swim-related network in the midbrain and the hindbrain^11,34^ (component 1), a network in the hindbrain that bias the fish’s swimming to the left side and to the right side^8,35^ (component 2, 3), and neurons in the optic tectum, the dorsal raphe and the thalamus that respond to visual feedback during motor learning task^13^ (component 4). These results demonstrate that our open-source PyZebrascope allows us to perform whole-brain neural activity imaging in behaving zebrafish in a quality comparable with the pioneering studies that relied on commercially developed software.

## Discussion

Here we described the development of the open-source Python software, PyZebrascope, that controls a light-sheet microscope designed for neural activity imaging experiments in zebrafish. Its intuitive graphical user interfaces and ease of managing complex device parameters allow the users to minimize efforts in setting up consistent whole-brain imaging experiments across sessions. The choice of Python as a programming language allowed us to implement an advanced microscope control algorithm, such as automatic focusing, which further reduces the tedious efforts for ensuring the quality of multi-beam volumetric scans. PyZebrascope and its components are written in a generalizable manner, enabling research teams that use different types of devices to adapt the code to their configurations.

PyZebrascope is available from Github (https://github.com/KawashimaLab/PyZebraScope_public). It only requires pre-installation of Anaconda Python package, μManager package^28^ with a matching version of its Python interface (pymmcore), a Python library for controlling data acquisition board (ni-daqmx), a Python library for GPU computing (CuPy), and a few other Python packages, all free of cost. Therefore, we expect that PyZebrascope will help disseminate the whole-brain imaging technique throughout the zebrafish neuroscience community.

The modular architecture of PyZebrascope further enables advanced microscope control and image processing algorithms. For example, a common issue during whole-brain imaging in zebrafish is the sample’s slow drift resulting from gravity force, tail motions of unparalyzed fish^14^, or the pressure of pipette attachment during fictive recording in paralyzed fish^11,13,30^. A small amount of drift, especially in the axial direction along which the volumetric scan is undersampled, can result in the loss of neurons during imaging because the neuronal diameter in the zebrafish brain is usually less than 5 μm. Given that such sample drift occurs at a slow rate, it can be occasionally calculated by using GPU-based algorithms during experiments to compensate for the drift by moving the 3-axis stage holding the sample. The effectiveness of such online drift correction is demonstrated in a multi-beam light-sheet microscope^36^. Therefore, implementing such an algorithm will increase the success rate of whole-brain imaging experiments in zebrafish, which are typically laborious and time-consuming.

It will also be possible to implement real-time image processing features to identify neurons of specific activity patterns for subsequent neural perturbation experiments at a single-cell resolution^19,37^. Real-time registration and segmentation of individual neural activity across the brain will require large computing resources and may not be feasible on the microscope control computer itself. Nonetheless, it will be possible to calculate, for example, trial-averaged neural activation maps based on a simple recurrent formula if the behavioral events or sensory stimulus is set to occur at regular volumetric scan intervals. Once we identify neurons that show specific activity patterns, we can further laser-ablate or photo-stimulate these populations by using an open-source Python resource for holographic two-photon stimulation^38^. Such experiments will allow us to investigate the functional roles of neurons that show specific activity patterns beyond what is obtainable by genetic labeling of neurons.

Lastly, PyZebrascope will enable further development of voltage imaging techniques in zebrafish. Voltage imaging requires high image resolution, high photon collection efficiency, and high imaging speed at the rate of at least 300 frames per second^22,39^. Light-sheet microscopy has the potential of realizing these conditions at a brain-wide scale in zebrafish. For example, it will be possible to implement a custom module that performs single-plane high-speed imaging across multiple Z planes in a sequential manner while the zebrafish perform a simple, stereotypical sensorimotor task as demonstrated for calcium imaging in zebrafish^30^. Implementing such a capability will advance our understanding of how whole-brain neural dynamics control animal behaviors at a millisecond timescale.

## Methods

### PyZebrascope

All the modules of PyZebrascope were written in Python programming language on Spyder IDE (https://www.spyder-ide.org/). We used a PC for microscope control that has Windows 10 operation system, two processors (Intel Xeon Gold 6244), 384 GB of DDR4 memory, a GPU (nVidia GeForce GTX1660), and a 15 TB SSD drive (Micron 9300 Pro, >3GB/s writing/reading speed). We followed manufacturers’ manuals for connecting devices, including the camera, data acquisition board and sample stages.

### Light-sheet microscope

We designed a custom light-sheet microscope that has a virtual reality setup for behavioral recordings, two optical paths for excitation beams, two cameras for fluorescence detection, and additional space for future implementation of optical paths for the third path for excitation beam and neural perturbations (**Fig. S1**). We designed this microscope using Inventor Professional software (Autodesk), and our CAD model is available upon request. We listed commercially available parts and custom-designed parts for our custom light-sheet microscope in **Supplementary Table 1**.

### Zebrafish experiments

We used a 6-day old transgenic zebrafish that pan-neuronally express nuclear-localized, genetically-encoded calcium indicators^33^ (Tg(HuC:H2B-GCaMP7f)^jf96^, courtesy of Dr. Misha Ahrens) for the imaging experiment. The zebrafish was immobilized and mounted to an imaging chamber as described previously^13^. Briefly, the fish larvae were immobilized by bath application of α-Bungarotoxin (B1601, Fisher Scientific, 1mg/ml) dissolved in external solution (in mM: 134 NaCl, 2.9 KCl, 2.1 CaCl2, 1.2 MgCl2, 10 HEPES, 10 glucose; pH 7.8; 290 mOsm) for 25-30 seconds and embedded in agarose on a custom-made pedestal inside a glass-walled chamber with a diffusive screen underneath the fish (**Fig. 1c**). Agarose around the head was removed with a microsurgical knife (#10318-14, Fine Science Tools) to minimize the scattering of the excitation laser. Laser power from the side beam path was, on average, approximately 21 µW. The distance between the fish and the display was about 4 mm.

We performed a fictive recording of fish’s swim patterns during a motor learning task^13^. Electric signals from motor neuron axons in the tail were recorded using borosilicate pipettes (TW150-3, World Precision Instruments) pulled by a horizontal puller (P-1000, Sutter) and fire-polished by a microforge (MF-900, Narishige). The pipettes were filled with fish-rearing water and connected to the tail using minimal negative pressure. Swim signals were recorded using an amplifier (RHD2132 amplifier connected to RHD-2000 interface board, Intan Technologies). We used custom-written Python software (available upon request) for executing the same algorithms for closed-loop virtual reality experiments as the software used in previous studies^13,22^. It samples signals from the above amplifier at 6 kilohertz, automatically detects swim events and moves the visual stimulus projected below the fish in a closed-loop with a delay of 35 milliseconds.

### Data analysis

We processed acquired imaging data on a Linux server in High Performance Computing (HPC) division in the Weizmann Institute of Science. This server has two Xeon processors (Xeon Gold 6248, Intel), 384 GB RAM, 13-TB SSD array, and a GPU computing board (Tesla V100, nVidia). We performed data processing by using custom Python scripts that execute the same algorithms as those established in our previous work^13^. All the analyses of imaging data were performed on a remote JupyterLab environment (https://jupyterlab.readthedocs.io/).

Briefly, we first registered time-series images from the same Z-planes by using phase correlation algorithms on the above GPU. We then examined residual drifts in the lateral and axial directions and discarded data with excessive drifts (>5 μm) in either direction. We then identified individual neurons that express nuclear-localized GCaMP based on the average image by using an algorithm for detecting circular shapes in images. We then extracted fluorescent time series from the central part of identified neurons (49 pixels). We identified 79,176 neurons across the brain in the experiment described in Fig. 4. We calculated the baseline fluorescence trace for each extracted fluorescence trace by taking the rolling percentile of the bottom 30% with a window size of 2 minutes and then divided the original fluorescent time series by this baseline trace to obtain ΔF/F time series for each neuron.

For the analyses shown in Fig. 4b, we focused on neurons whose responses were reliable across multiple task trials. For extracting such neurons, we created a matrix of the average ΔF/F in each task period (12 periods during motor learning task^13^) in every trial and performed one-way ANOVA across trials for individual neurons. In this way, we identified 36, 818 neurons with significant p-values (p<0.001) for response reliability. We then extracted neurons for each brain area by their spatial locations. We classified neurons into three groups in each area by applying a k-means clustering method (n=3) to the trial average activities of neurons. We then picked a neuron that shows the largest response amplitude from each identified cluster in each brain area and plotted their time series in Fig. 4b.

For the analysis shown in Fig. 4c, we used the same set of 36,818 neurons identified in the above statistical test for response reliability. We normalized the ΔF/F trace of each neuron by its 99-percentile value and clipped values more than 1. We then applied non-negative matrix factorization to the activity of neurons (n=4 components). We extracted neurons whose weight is more than 0.2 for each component and color-plotted their locations in the top projection and side projection images of the brain.

## Author contribution

T.K. and R.B. conceived the project. R.B. established the device control and first versions of PyZebrascope. T.K. designed the custom light-sheet microscope, organized the later versions of PyZebrascope and conducted data analyses. R.H. provided initial user feedback for refining PyZebrascope and collected whole-brain imaging data in behaving zebrafish. M.N.K., and K.H. developed algorithms for automatic image focusing and contributed a figure. T.K. wrote the manuscript with inputs from all authors.

## Acknowledgment

We thank Dorel Malamud and other members of Kawashima laboratories for experimental help, members of Veterinary Resource at Weizmann Institute of Science for animal care, Meir Alon, Haim Sade, Alex Jahanfard and members of Physics Core Facility at Weizmann Institute of Science for machining and assembling custom microscope parts, and Uri Rosen for the maintenance of our computing server in the High Performance Computing unit. We thank Igor Negrashov, Davis Bennet, Raghav Chhetri, Nikita Vladimirov and Steven Sawtelle (HHMI Janelia) for their advice on our initial design of the microscope components and PyZebrascope, and Niels Cautaerts (Data Minded) for his advice on GPU-based codes. We thank Inbal Shainer for her critical reading of the manuscript. M.N.K. is supported by iNAMES MDC-Weizmann Research School, “Imaging from NAno to MESo”. This research is supported by Azrieli faculty fellowship (T.K.), Dan Lebas & Ruth Sonnewend (T.K.), Birnbach Family foundation (T.K.), Internal grant from the Center for New Scientists at Weizmann Institute of Science (T.K.), Helmholtz Imaging (K.H.), and an Oak Ridge National Laboratory, Laboratory Directed Research and Development Strategic Hire grant (K.H).

---

**Supplementary Figure 1:**
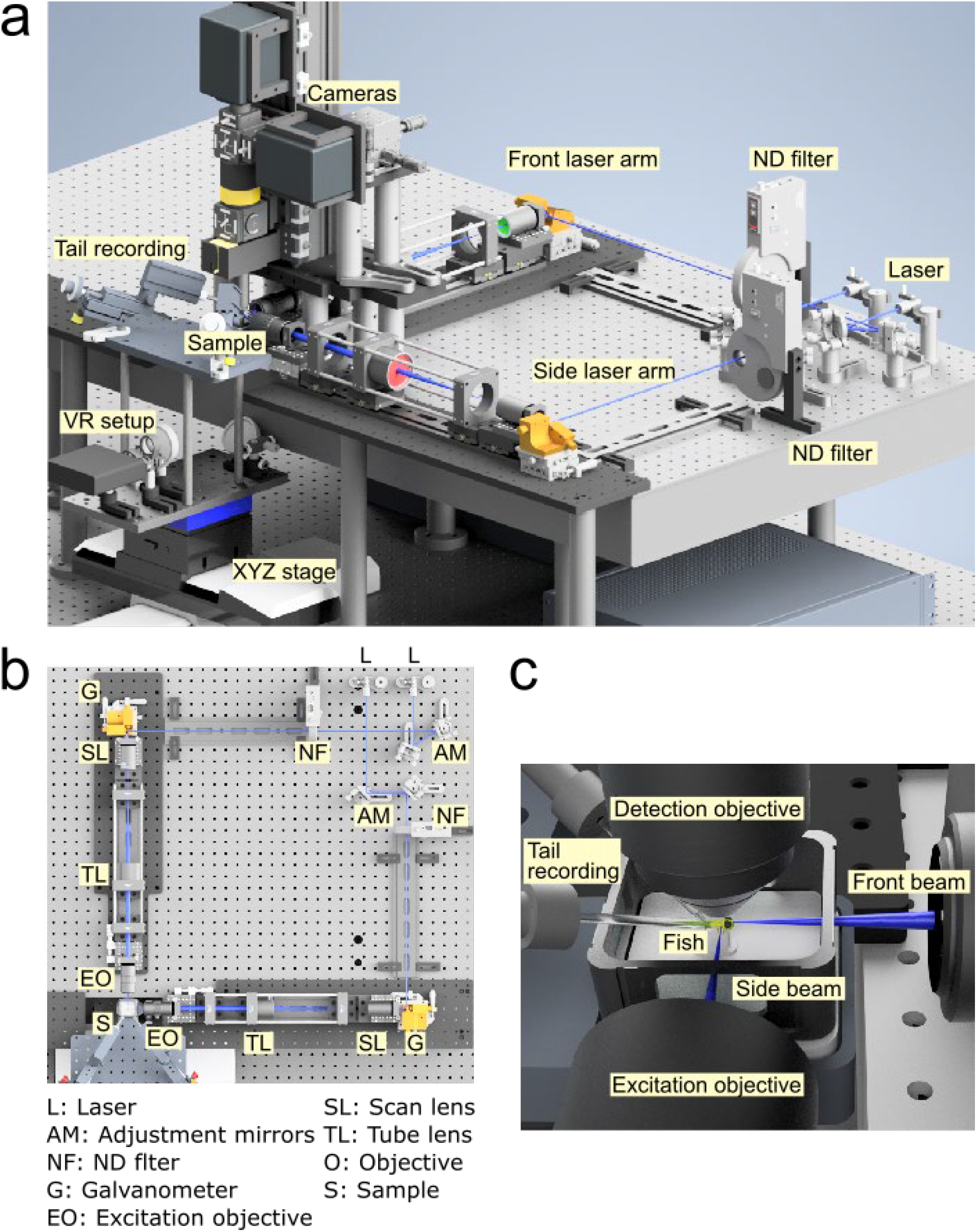
3D rendering of our custom lightsheet microscope. **(a)** A rendered view of the microscope. **(b)** Optical path of the excitation beams. **(c)** Illumination of two excitation beams in the brain of zebrafish in a water chamber.

**Supplementary Figure 2:**
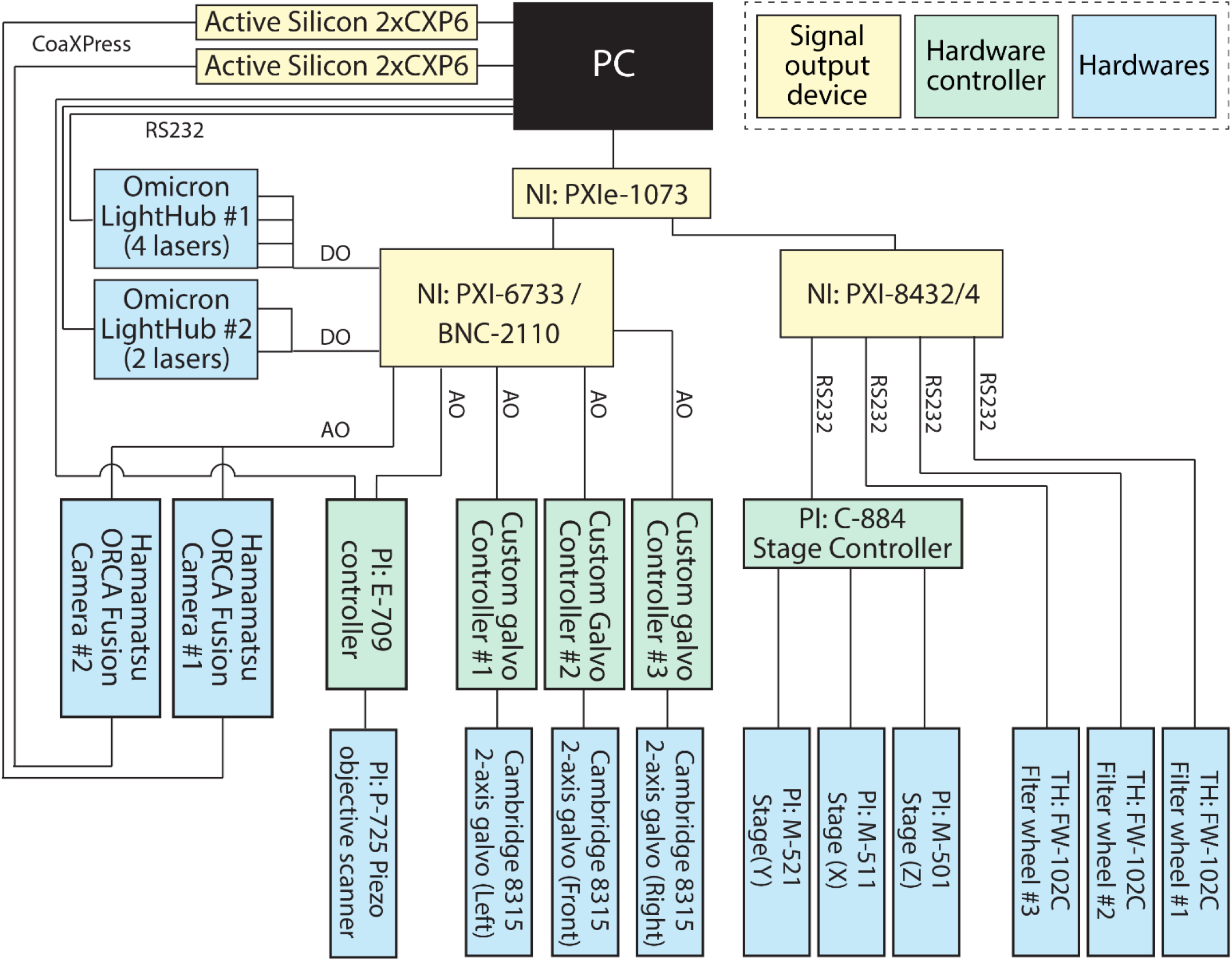
Wiring diagram of a custom lightsheet microscope. Wiring diagrams among signal input/output devices (yellow), hardware controller (green) and microscope hardware (green). CoaXPress, camera connection via CXP6 coaxial cables; AO, analog outputs via BNC coaxial cables; DO, digital outputs; RS232, communication via DB9 serial cables.

**Supplementary Table 1:** List of major parts for light-sheet microscope.

| Item | Details | Supplier |
| --- | --- | --- |
| PC |  |  |
| Windows 10 PC | Two Xeon 6244, 192 GB RAM, 15 TB SSD, 10GbEconnection | Access Technology ltd, Israel |
| Optical table |  |  |
| Custom optical table | Custom RPR table Top 4.0' x 7.0' x 8.0" | (Manufacturer)<br>Newport, USA<br>(Local supplier)<br>New technology, Israel |
| Vibration isolator | S-2000A-422 |  |
| Optical breadboard | Custom breadboard 2.5' x 5.0' x 4.32" |  |
| Optical support rod | XP-10 |  |
| DAQ |  |  |
| PXI chassis | NI PXIe-1073 | (Manufacturer)<br>National Instruments, USA<br>(Local rep)<br>STEM-Innovation, Israel |
| Analog outputs | NI PXIe-6341 |  |
| Analog inputs | NI PXI-6733 |  |
| Serial inputs/outputs | NI PXI-8432/4 |  |
| Connector block | NI BNC-2110 |  |
| Camera |  |  |
| sCMOS camera | ORCA Fusion BT | Hamamatsu Photonics Israel ltd, Israel |
| Frame grabber | Active Silicon 2xCXP6-2PE8 |  |
| Laser |  |  |
| Laser combiner #1 (four wavelengths) | LIGHTHUB+ laser engine (488, 532,561 and 594 nm) | Omicron-Laserage Laserprodukte GmbH, Germany |
| Laser combiner #2 (two wavelengths) | LIGHTHUB+ laser engine (488,561 nm) |  |
| Neutral density filter | NDQ-100, 150, 200, 300, 400 | CVI laser optics, USA |
| Laser goggles | Cat # : DYE, GRY, ZS2 | NoIR Laser Company, USA |
| Scanning unit |  |  |
| 2-axis galvanometer | 6SD11872 kit (8315K galvanometers with silver-coated 5 mm mirrors, single axis servo drivers) | (Manufacturer)<br>Cambridge Technology, USA<br>(Local supplier)<br>New technology, Israel |
| Left-handed XY mount | 61021505L20 |  |
| 2-axis manual translation stage | TSD-602S / SD-602SR | OptoSigma Europe SAS, France |
| Custom mount | Aluminum mount between manual stage and galvanometer mount | Physics Core facility at Weizmann Institute. |
| Custom linear power supply | Linear 28V power supply for two-axis galvanometer (www.janelia.org/open-science/galvo-driver). |  |
| Sample Stage |  |  |
| 3-axis stage | M-521.DD1 (Y)<br>M-511.DD1 (X)<br>M-501.DG1 (Z)<br>C-844.4DC (Controller) | (Manufacturer)<br>Physik Instrumente, Germany<br>(Local supplier)<br>GOA-Tech, Israel |
| Projector | Optoma LV130 | Videoset tech, ltd. Israel |
| Wratten filter #25 (red) | Cat # 53699 | (Manufacturer) Edmund Optics, USA<br>(Local supplier) Prolog Optics, Israel |
| Custom chamber for zebrafish | Delrin chamber, acrylic fish pedestal, acrylic plate, polycarbonate base | Physics Core facility at Weizmann Institute. |
| Custom stage for fish chamber | Anodized aluminum |  |
| Electrophysiology setup |  |  |
| 3-axis manual manipulator, clamp, headstage mount | MX130R, MX130L, MXB-3h, MXC | (Manufacturer)<br>Siskiyou Corporation, USA<br>(Local supplier)<br>Lahat Technologies, Israel |
| Electrode holder | Axon HL-U | Molecular Devices, USA |
| Amplifier and interface board | RHD2000, RHD2132 | Intan Technologies, USA |
| 1 mm pins, jacks, sleeves | 08F150, 20F2610, 08F162 | Bürklin Elektronik, Germany |
| Custom headstage | 3D printed, PLA |  |
| Excitation and detection optics |  |  |
| Breadboard for excitation optics | MB636 | (Manufacturer)<br>Thorlabs Inc, USA<br>(Local supplier)<br>Rosh electoptics, Israel |
| Damped post under galvanometer | DP14A |  |
| ND filter wheel | FW102C |  |
| Tube lens for detection objective | TTL200-A |  |
| Scan lens | SL50-CLS2 |  |
| Tube lens | TL200-CLS2 |  |
| Excitation objective | MY5X-802 |  |
| 1-axis stage | LNR25D |  |
| Filter, dichroic | GFP filter (FF03-525/50-25)<br>RFP filter (FF01-607/70-25) | (Manufacturer)<br>Semrock, USA<br>(Local supplier)<br>Lahat Technologies, Israel |
| Detection objective | CFI75 LWD 16X W | (Manufacturer)<br>Nikon, Japan<br>(Local supplier)<br>Agentek, Israel |
| Piezoelectric drive for the detection objective | P-725.4CA (400 μm actuator)<br>E-709 Controller | (Manufacturer)<br>Physik Instrumente GmbH, Germany<br>(Local supplier)<br>GOA-Tech Ltd, Israel |
| Miscellaneous |  |  |
| Stereomicroscope | Microscope (SM745B-V203)<br>Flexible arm (ASC) | AmScope, USA |
| Custom mount for detection optics | Aluminum | Physics Core facility at Weizmann Institute. |
| Miscellaneous custom parts |  |  |
| Custom laser protection panels | Anodized aluminum sheet |  |
| Optomechanical parts | Mirrors, lens, mounts, rails, cages, rods, tubes, screws | (Manufacturer)<br>Thorlabs Inc, USA<br>(Local supplier)<br>Rosh electoptics, Israel |
| CAD model |  |  |
| The full CAD model is available upon request. |  |  |

## References

1. Ahrens, M. B. & Engert, F. Large-scale imaging in small brains. Curr. Opin. Neurobiol. 32, 78–86 (2015).

2. Chen, T.-W. et al. Ultrasensitive fluorescent proteins for imaging neuronal activity. Nature 499, 295–300 (2013).

3. Inoue, M. et al. Rational Engineering of XCaMPs, a Multicolor GECI Suite for In Vivo Imaging of Complex Brain Circuit Dynamics. Cell 177, 1346–1360.e24 (2019).

4. Keller, P. J., Schmidt, A. D., Wittbrodt, J. & Stelzer, E. H. K. Reconstruction of zebrafish early embryonic development by scanned light sheet microscopy. Science (80-.). 322, 1065–1069 (2008).

5. Sofroniew, N. J., Flickinger, D., King, J. & Svoboda, K. A large field of view two-photon mesoscope with subcellular resolution for in vivo imaging. Elife 5, (2016).

6. Demas, J. et al. High-speed, cortex-wide volumetric recording of neuroactivity at cellular resolution using light beads microscopy. Nat. Methods 2021 189 18, 1103–1111 (2021).

7. Voleti, V. et al. Real-time volumetric microscopy of in vivo dynamics and large-scale samples with SCAPE 2.0. Nat. Methods 2019 1610 16, 1054–1062 (2019).

8. Ahrens, M. B., Orger, M. B., Robson, D. N., Li, J. M. & Keller, P. J. Whole-brain functional imaging at cellular resolution using light-sheet microscopy. Nat Methods 10, 413–420 (2013).

9. Panier, T. et al. Fast functional imaging of multiple brain regions in intact zebrafish larvae using Selective Plane Illumination Microscopy. Front. Neural Circuits 7, 65 (2013).

10. Freeman, J. et al. Mapping brain activity at scale with cluster computing. Nat. Methods 11, 941–950 (2014).

11. Vladimirov, N. et al. Light-sheet functional imaging in fictively behaving zebrafish. Nat. Methods 11, 883–884 (2014).

12. Chen, X. et al. Brain-wide Organization of Neuronal Activity and Convergent Sensorimotor Transformations in Larval Zebrafish. Neuron 100, 876–890.e5 (2018).

13. Kawashima, T. et al. The Serotonergic System Tracks the Outcomes of Actions to Mediate Short-Term Motor Learning. Cell 167, 933–946.e20 (2016).

14. Markov, D. A., Petrucco, L., Kist, A. M. & Portugues, R. A cerebellar internal model calibrates a feedback controller involved in sensorimotor control. Nat. Commun. 2021 121 12, 1–21 (2021).

15. Mu, Y. et al. Glia Accumulate Evidence that Actions Are Futile and Suppress Unsuccessful Behavior. Cell 178, 27–43.e19 (2019).

16. Mancienne, T. et al. Contributions of Luminance and Motion to Visual Escape and Habituation in Larval Zebrafish. Front. Neural Circuits 15, 115 (2021).

17. Migault, G. et al. Whole-Brain Calcium Imaging during Physiological Vestibular Stimulation in Larval Zebrafish. Curr. Biol. 28, 3723–3735.e6 (2018).

18. Adriá Ponce-Alvarez, A. et al. Whole-Brain Neuronal Activity Displays Crackling Noise Dynamics Article Whole-Brain Neuronal Activity Displays Crackling Noise Dynamics. Neuron 100, (2018).

19. Vladimirov, N. et al. Brain-wide circuit interrogation at the cellular level guided by online analysis of neuronal function. Nat. Methods 15, 1117–1125 (2018).

20. Yang, C. et al. All-optical imaging and manipulation of whole-brain neuronal activities in behaving larval zebrafish. Biomed. Opt. Express, Vol. 9, Issue 12, pp. 6154-6169 9, 6154–6169 (2018).

21. Burgstaller, J. et al. Light-sheet imaging and graph analysis of antidepressant action in the larval zebrafish brain network. bioRxiv 618843 (2019). doi:10.1101/618843

22. Abdelfattah, A. S. et al. Bright and photostable chemigenetic indicators for extended in vivo voltage imaging. Science 365, 699–704 (2019).

23. Böhm, U. L. et al. Voltage imaging identifies spinal circuits that modulate locomotor adaptation in zebrafish. Neuron 0, (2022).

24. Pologruto, T. A., Sabatini, B. L. & Svoboda, K. ScanImage: Flexible software for operating laser scanning microscopes. Biomed. Eng. Online 2, 1–9 (2003).

25. Vladimirov, N. mesoSPIM-control: Image acquisition software for mesoSPIM light-sheet microscopes. Available at: https://github.com/mesoSPIM/mesoSPIM-control.

26. Harris, C. R. et al. Array programming with NumPy. Nat. 2020 5857825 585, 357–362 (2020).

27. Van Der Walt, S. et al. Scikit-image: Image processing in python. PeerJ 2014, e453 (2014).

28. Edelstein, A. D. et al. Advanced methods of microscope control using μManager software. J. Biol. Methods 1, 10 (2014).

29. Favre-Bulle, I. A., Vanwalleghem, G., Taylor, M. A., Rubinsztein-Dunlop, H. & Scott, E. K. Cellular-Resolution Imaging of Vestibular Processing across the Larval Zebrafish Brain. Curr. Biol. 28, 3711–3722.e3 (2018).

30. Ahrens, M. B. et al. Brain-wide neuronal dynamics during motor adaptation in zebrafish. Nature 485, 471–7 (2012).

31. Saska, D., Pichler, P., Qian, C., Buckley, C. L. & Lagnado, L. μSPIM Toolset: A software platform for selective plane illumination microscopy. J. Neurosci. Methods 347, 108952 (2021).

32. Mizutani, R. et al. A method for estimating spatial resolution of real image in the Fourier domain. J. Microsc. 261, 57–66 (2016).

33. Dana, H. et al. High-performance calcium sensors for imaging activity in neuronal populations and microcompartments. Nat. Methods 16, 649–657 (2019).

34. Orger, M. B., Kampff, A. R., Severi, K. E., Bollmann, J. H. & Engert, F. Control of visually guided behavior by distinct populations of spinal projection neurons. Nat. Neurosci. 11, 327–33 (2008).

35. Dunn, T. W. et al. Neural Circuits Underlying Visually Evoked Escapes in Larval Zebrafish. Neuron 89, 613–628 (2016).

36. Royer, L. A. et al. ClearVolume: open-source live 3D visualization for light-sheet microscopy. Nat. Methods 2015 126 12, 480–481 (2015).

37. dal Maschio, M., Donovan, J. C., Helmbrecht, T. O. & Baier, H. Linking Neurons to Network Function and Behavior by Two-Photon Holographic Optogenetics and Volumetric Imaging. Neuron 94, 774–789.e5 (2017).

38. Pozzi, P. & Mapelli, J. Real Time Generation of Three Dimensional Patterns for Multiphoton Stimulation. Front. Cell. Neurosci. 15, 34 (2021).

39. Cohen, A. E. & Venkatachalam, V. Bringing Bioelectricity to Light. (2014). doi:10.1146/annurev-biophys-051013-022717

